# Condensin controls mitotic chromosome stiffness and stability without forming a structurally contiguous scaffold

**DOI:** 10.1101/384982

**Authors:** Mingxuan Sun, Ronald Biggs, Jessica Hornick, John F. Marko

## Abstract

During cell division, chromosomes must be folded into their compact mitotic form to ensure their segregation. This process is thought to be largely controlled by the action of condensin SMC protein complexes on chromatin fibers. However, how condensins organize metaphase chromosomes is not understood. We have combined micromanipulation of single human mitotic chromosomes, sub-nanonewton force measurement, siRNA interference of condensin subunit expression, and fluorescence microscopy, to analyze the role of condensin in large-scale chromosome organization. Condensin depletion leads to a dramatic (~10 fold) reduction in chromosome elastic stiffness relative to the native, non-depleted case. We also find that prolonged metaphase stalling of cells leads to overloading of chromosomes with condensin, with abnormally high chromosome stiffness. These results demonstrate that condensin is a main element controlling the stiffness of mitotic chromosomes. Isolated, slightly stretched chromosomes display a discontinuous condensing staining pattern, suggesting that condensins organize mitotic chromosomes by forming isolated compaction centers that do not form a continuous scaffold.

## Introduction

Formation of mechanically stable compacted mitotic chromosomes is essential for faithful DNA segregation during cell division in eukaryotes. However, the molecular mechanisms underlying this process are still poorly understood. Condensins are multi-subunit protein complexes which have been implicated as key players in assembly and maintenance of mitotic chromosomes. Condensin’s function was initially illuminated through two types of experiments: antibody depletion of condensin from *Xenopus* egg extracts blocked *in vitro* chromosome assembly (Hirano and Mitchison, 1994), while mutation of condensin in yeast led to an increase in distance between pairs of chromosome loci (Strunnikov et al., 1995). Subsequently, condensin was demonstrated to be required for chromosome segregation: condensin-defective cells have chromosome segregation defects, with “anaphase bridges” formed between sister chromatids (Saka et al., 1994, Bhat et al., 1996, Sutani et al., 1999, Steffensen et al., 2001).

Metazoan cells possess two types of condensins: condensin I and II (Yeong et al., 2003, Ono et al., 2003). Together, they contribute to a wide variety of aspects of higher-order chromosome structure and dynamics. Condensin II is present in the nucleus during interphase, and participates in the early stage of mitotic chromosome folding during prophase. By contrast, condensin I is found in the cytoplasm, associates with chromosomes only after nuclear envelope breakdown (NEB), and is thought to establish an additional level of metaphase chromosome compaction. Condensin I and II share the same core subunits (SMC2 and SMC4, sometimes called CAP-E and CAP-C, respectively) which belong to the Structural Maintenance of Chromosomes (SMC) protein family, while having a distinct set of non-SMC subunits, including a kleisin subunit and two HEAT-repeat proteins: CAP-
H, CAP-G, and CAP-D2 for condensin I, and CAP-H2, CAP-G2, and CAP-D3 for condensin II (Hirano, 2006, Ono et al., 2003).

Depletion of condensins has been show to result in “fuzzy” chromosomes in organisms from *Drosophila* to human (Somma et al., 2003, Hudson et al., 2003, Ono et al., 2003), with recent experiments showing that metaphase chromosome structure is particularly sensitive to depletion of condensin II (Ono et al., 2017). It has been established that condensin I and II are differentially distributed along mitotic chromatids (Ono et al., 2003). These conclusions were mainly reached from imaging studies of fixed cells or using chromosome reconstitution in cell extracts. It is still unclear how condensins globally organize mitotic chromosome structure and quantitatively contribute to chromosome mechanical stiffness and stability, although a few studies have shown that metaphase chromosomes depleted in condensin have a more swollen conformation and are more easily deformed by hydrodynamic shear flow than unperturbed chromosomes (Green et al., 2012, Hudson et al., 2003, Ono et al., 2017).

Despite a variety of proposals of folding schemes for metaphase chromosome folding [e.g., axial scaffold (Adolphs et al., 1977, Earnshaw and Laemmli, 1983), helical loop (Bak et al., 1977), and chromatin network (Poirier and Marko, 2002) models], experiments have not indicated clearly which of these, if any, is most correct, although recent Hi-C studies do suggest a helical loop-array organization (Gibcus et al., 2018). Prior studies have also not clearly established whether condensin is organized into a continuous or discontinuous distribution along chromosome arms, although a recent super-resolution study does suggest the latter (Walther et al., 2018).

To examine the role of condensin in chromosome organization and mechanics, we carried out a study of isolated and micromanipulated human (HeLa) metaphase chromosomes, combining use of force-calibrated micropipettes, siRNA depletion of condensin subunits, and immunofluorescence analysis of condensin distributions using microspraying of antibodies on native unfixed chromosomes. This approach allows a quantitative measurement of structural changes induced in mitotic chromosomes by condensin depletion in human HeLa cells, as well as visualization of the condensin distribution along the chromosome arms from unperturbed cells. A key feature of our approach is the ability to image the condensin distribution on slightly stretched chromosomes, which allows us to determine the degree of connectivity of the condensin distribution along a single chromosome.

## Materials and Methods Cell culture

### cell cluture

Sample dishes were prepared using #1 microscope glass onto which 25 mm-diameter rubber O-rings were affixed using paraffin. HeLa cells were cultured in DMEM with 10% FBS (Gibco), 100U/ml penicillin/streptomycin (Fisher Scientific) in a 37°C incubator supplemented with 5% CO2.Experiments were performed on ~70% confluent samples. Metaphase cells were identified by phase-contrast imaging. For mitotic arrest experiments, cells were treated with 2.5 ng/μl colchicine (Sigma-Aldrich) or 1 ng/μl nocodazole (Sigma-Aldrich) for 14-18 hours.

### Microscopy and chromosome micromanipulation

Pipette micromanipulation and imaging were done on the stage of an inverted microscope (IX70; Olympus) using a 60X 1.4 NA oil immersion objective at 30 °C with a temperature controlled heater attached to the objective (Fig. 1A). Imaging was done using a CCD camera (Pelco, DSP B&W) with images acquired a frame grabber (IMAQ PCI-1408, National Instruments) controlled by homebuilt Labview (National Instruments) codes, or by a EMCCD camera (Ixon3, Andor) controlled by MetaMorph software (MetaMorph Inc.).

**Figure 1.**
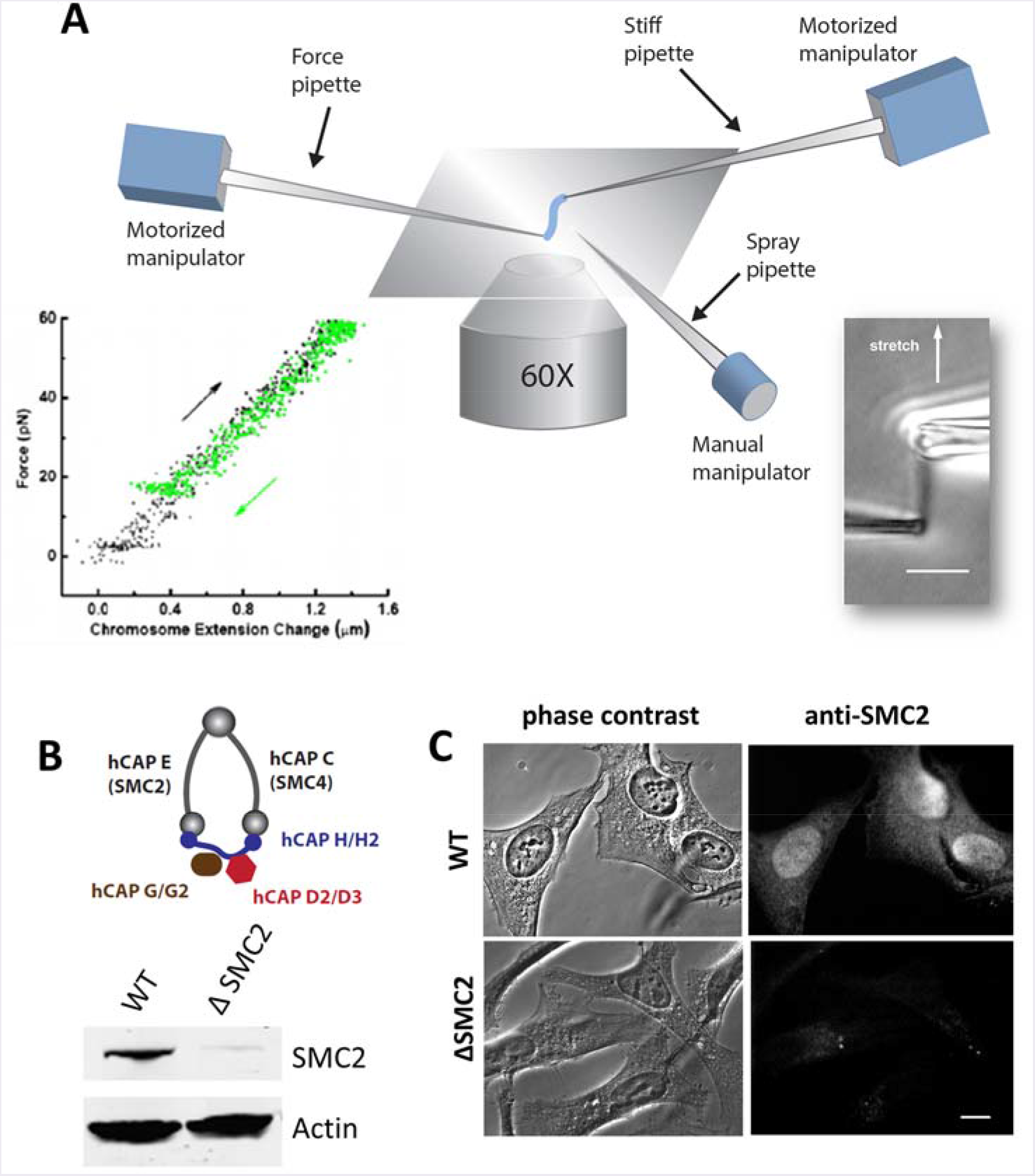
Combination of micromechanics measurement of mitotic chromosomes with RNAi depletion of condensin. (A) Schematic diagram of experimental micropipette setup for chromosome micromanipulation. One of the two pipettes (force pipette) has a long taper with a spring constant of 30-200 pN/μm, which is used to measure forces in the range of 10-1000 pN by monitoring its bending. A third pipette is used as the spray pipette to spray reagents directly onto an isolated chromosome. Lower right insert shows a mitotic chromosome from human HeLa cells, and the lower left insert shows its linear, reversible elasticity. (B-C) Depletion of SMC2 (hCAP-E) using siRNA in HeLa cells. Protein knockdown is verified by western blot (B), and (C) immunofluorescence staining of SMC2 in fixed cells 36 hrs after siRNA treatment, using anti-SMC2 antibody in both cases. Bar = 5 μm.

In brief, prometaphase cells were identified by phase-contrast imaging, and a micropipette filled with 0.05% v/v Triton X-100 (Fisher Scientific) in PBS pipette two micropipettes were positioned into the sample dish controlled by a motorized manipulator (MP-285; Sutter Instrument Co.) to destabilize the cell membrane by microspraying. After the chromosomes were released from the cell, two micropipettes were positioned into the sample dish. One of the two pipettes was pulled with a rather short taper so it was stiff, and the other one was pulled with a long taper so as to have a softer tip with spring constant of 30-200 pN/μm. The stiff pipette was used to catch one end of a single chromosome using aspiration. Then the floppy pipette was attached to the other chromosome end.

For micromanipulation, the stiff pipette was moved at a rate of ~0.008-0.040 μm/sec, slow enough to avoid the viscoelastic effects (Poirier et al., 2000). Bending of the force pipette was recorded to monitor the force applied on chromosomes. Each extension-relaxation measurement was repeated at least 3 times to ensure its reproducibility. Micromechanical data were collected using image analysis software written in Labview. Offline image analysis (line scan analysis, feature size and spacing analysis) was done using Fiji/ImageJ software (Schindelin et al., 2012).

### siRNA

Transfections were carried out by incubating 100 nM siRNA with transfection reagent Dharmafect I (Dharmacon) in DMEM. After 6 hours media were exchanged to fresh DMEM. Control experiments used non-targeting control RNA (Dharmacon, siGENOME Non-Targeting siRNA #1, 5’-UGGUUUACAUGUCGACUAA UU-3’), or water, as indicated. Targeting RNAs used were: SMC2/Cap-E sequence: #1: 5’-UGCUAUCACUGGCUUAAAUTT-3’, sequence #2: 5’-CAUAUUGGACUCCAUCUGCTT-3’; hCAP-G: sequence #1: 5’-
UCAGAUAUGGAAGAUGAUGTT-3’, sequence #2: 5’-GUC UCAUGAAGCAAACAGCTT-3 ‘; hCAP-G2: sequence #1: 5’-UGAUUG CAUCCAGGACUUCTT-3’, sequence #2: 5’-UAGCAAAGCUGACACGUTT-3’.

### Antibodies

The following primary antibodies were used: rabbit polyclonal antibody against SMC2/hCAP-E (07-710, Upstate), mouse monoclonal antibody against hCAP-G (H00064151-M01, Novus Bio), and rabbit polyclonal antibody against hCAP-G2 (NB100-1813, Novus Bio). Secondary antibodies used for immunofluorescence were: Alexa 488 donkey anti-mouse IgG (Invitrogen), Alexa 594 goat anti-mouse IgG (Invitrogen), Alexa 488 goat anti-rabbit IgG (Invitrogen), and Alexa 594 donkey anti-rabbit IgG (Invitrogen). Centromere labeling was done using a labeled primary antibody (Texas red-conjugated CREST antibody, 15-235-T, Antibodies Inc). Secondary antibodies used in the western blots were: IRDye 800CW goat anti-mouse IgG, IRDye 680LT labeled goat anti-rabbit (Li-Cor).

Primary Fab fragment generation for anti-SMC2 was done using Papain digestion (Pierce, Fab Fragmentation Kit 44685) with fragmentation verified by SDS-PAGE. Secondary Fab fragments were purchased (Alexa 594-labeled goat-anti-rabbit Fab fragment, 111-587-003 Jackson ImmunoResearch).

### Immunofluorescence staining

For whole-cell experiments, cells on cover glass were washed with PBS, and then fixed by 4% (w/v) formaldehyde in PBS solution for 20 mins at room temperature, then washed with PBS three times. Fixed cells were permeabilized with 0.2% (w/v) Triton X-100 in PBS, incubated with Odyssey blocking buffer (Li-Corr) for 1 hour, and then incubated with primary antibodies overnight at 4C at a dilution of 1:1000 (stock antibody concentration ~1 mg/ml; dilution ~1 μg/ml in PBS). After primary incubation, the slides were washed with 3 times in PBS. Secondary treatment was done at a dilution of 1:500 for 1 hour at room temperature. Three more washes were then performed with PBS.

For single-chromosome microspraying, primary and secondary antibodies were diluted in 50% PBS with 0.5% Casein; primaries were diluted to 10 μg/ml and secondaries to 20 μg/ml. Antibody solutions were loaded into the spray pipettes, pulled and cut to have a ~5 μm opening. The spray pipettes were mounted on a three-axis manual manipulator (Taurus, World Precision Instruments), and positioned manually ~50 μm away from and pointing towards the isolated chromosome. The microspray was carried out for 10 mins with applied pressure of 10-100 Pa, and stopped for several minutes to allow spray solution (total volume of ~10 ul) to diffuse away into the 1.8 ml sample dish (Poirier and Marko, 2002, Pope et al., 2006). After single chromosome extraction, chromosomes were sprayed with blocking solution (PBS with 0.5% Casein), primary antibody and secondary antibody sequentially. Images were acquired using

Experiments with Fab fragments were carried out according to the same protocol as for full-length antibodies, except primary and secondary Fab fragments were diluted to 20 μg/ml.

## Results

### Morphological changes in human mitotic chromosomes following siRNA depletion of condensin

To correlate condensin level with chromosome morphology and chromosome mechanics, we combined micropipette based chromosome isolation with siRNA depletion of condensin SMC2 core subunits in HeLa cells (Fig. 1). We note that SMC2 depletion disrupts both condensin I and II complexes. In brief, HeLa cells were transfected with 50-100 nM of either non-targeting control siRNA or siRNA targeting the condensin SMC2 subunit (Dharmacon). The condensin protein knockdown level was ~90% after 36 hours following siRNA treatment, as quantified by Western blots (Fig. 1B, Fig. S1), and immunofluorescence staining (Fig. 1C). No depletion of SMC2 was observed in cells treated with the non-targeting control siRNA (Fig. S1).

Following siRNA treatment, metaphase cells were identified, and a spray pipette was introduced into the cell culture to gently disturb the cell membrane using dilute Triton X-100 solution, releasing metaphase chromosomes from the cell (Sun et al., 2011). This allowed a second micropipette to be used to capture either the whole genome (the full set of metaphase chromosomes from a cell), or individual chromosomes (Fig. S2), via gentle aspiration. We note that the set of metaphase chromosomes in a single genome are generally attached to one another by interchromosome linker filaments (Hoskins, 1968, Maniotis et al., 1997); these fibers can be broken to isolate a single chromosome. In our experiments, whole genomes or individual chromosomes were mechanically analyzed and imaged immediately in their native state in normal cell culture medium, without the need for cell-cycle synchronization or fixation.

We examined chromosome morphological change at times ranging from 24 to 72 hours after SMC2 siRNA treatment, by examination of whole genomes extracted from cells. Compared with chromosomes from non-siRNA-treated cells (referred to below as wild-type or WT), which appear as homogenous rod-like structures (Fig. 2Aa), we observed progressive morphological changes of chromosomes in condensin-depleted cells (Fig. 2A, b-d). Chromosomes at 24 hours after siRNA treatment began to show fragmented structures, with thinner and thicker regions apparent within single chromosomes. 48 hours after siRNA treatment, chromosomes appeared fluffy or frayed. Eventually, after 72 hours, the whole genome appeared to be a mesh-like mass of chromatin, poorly compacted without visible signs of individual chromosomes, despite having been released from the nucleus following NEB.

**Figure 2.**
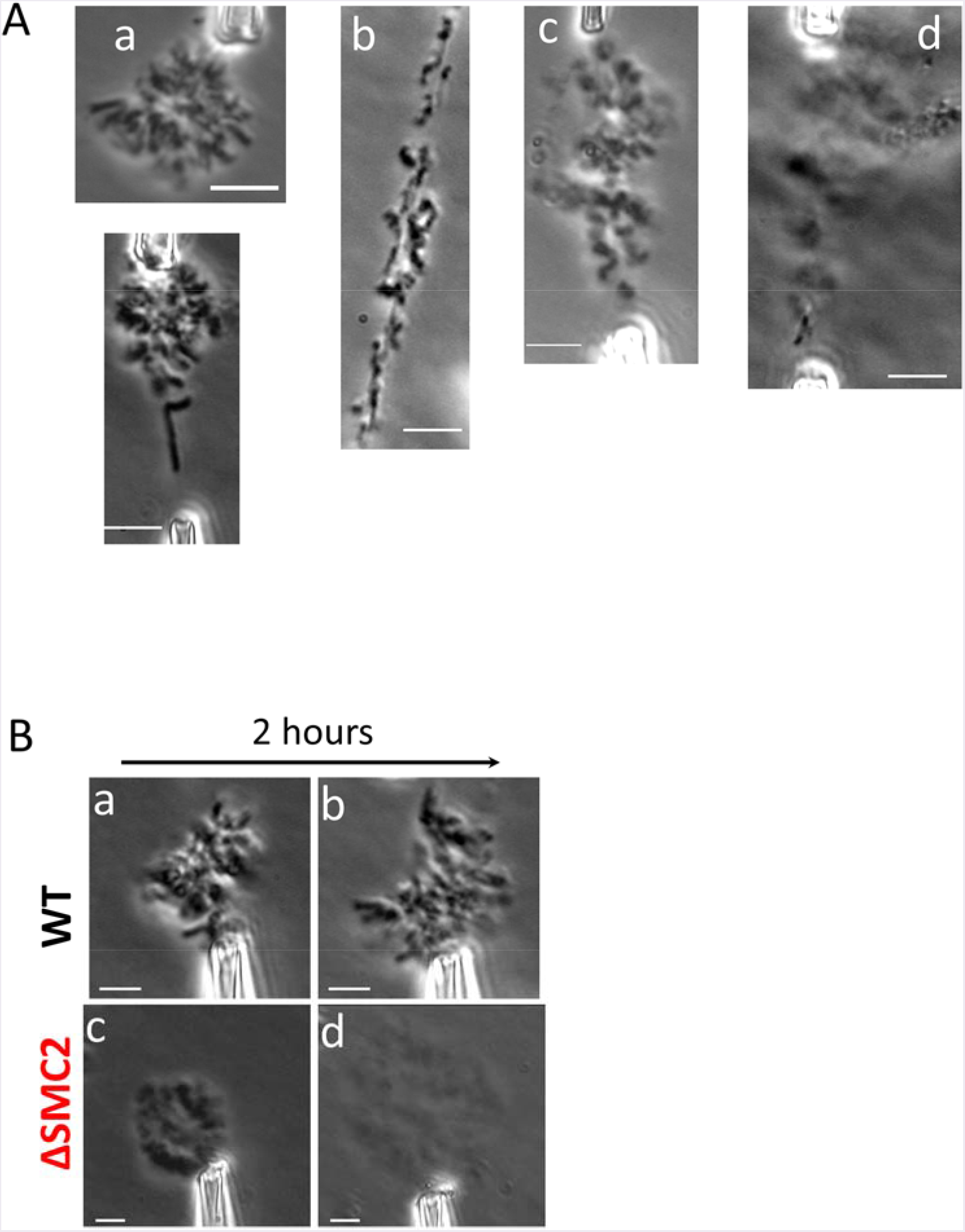
siRNA mediated depletion of SMC2 results in progressive change in chromosome morphology, and affects genome stability. (A) Whole genome isolated with micropipettes from WT (untreated) HeLa cells (a), and siRNA treated cells at varied times following siRNA treatment ((b), 24 hr; (c), 48 hr; (d), 72 hr). After opening up the cell membrane, chromosomes were taken out of the cells using aspiration, and imaged immediately in the cell culture medium (DMEM). (B) Representative examples of genomes isolated from control cells ((a) and (b), top row) and from cells 36 hours after treatment by SMC2-targeting siRNA ((c) and (d), bottom row). Phase contrast images were taken either right after chromosome isolation (left column) or after 2 hours sitting in cell culture medium (right column). Only one micropipette was used to hold any given isolated whole genome. Bar = 5μm.

### Condensin controls the stability of mitotic chromosomes

WT metaphase chromosomes are very stable when isolated to the cell culture medium (Pope et al., 2006). Fig. 2Ba shows a phase contrast image of a WT metaphase genome held by one micropipette immediately after having been isolated from a cell. Over two hours of exposure to cell culture media, no significant change in chromosome structure was observed (Fig. 2Bb). Since the chromosomes in a genome are connected together by interchromosome linkers, the chromosomes stayed clustered together and did not diffuse away from one another, even when only one part of the genome was held by one micropipette.

For condensin-depleted (SMC2 siRNA treated) cells, the isolated genome appeared poorly compacted immediately after extraction. Fig. 2Bc shows a whole genome isolated from cells 48 hours after siRNA treatment, held by a micropipette. Furthermore, after two hours outside the cell condensin-depleted chromosomes spontaneously dissolved, expanding more than three times in area, and becoming much lighter under phase contrast (Fig. 2Bd). The size of the genome was quantified using averaged line scans of images collected from phase-contrast imaging. The genome size was determined by measurement of the width of the phase contrast intensity distribution at half-maximum, relative to background intensity (isolation of the genome far from the cells and cell debris allows a well-defined background measurement). We found a 3.5 ± 0.5 times increase in area; the volume increase is likely to be even larger due to expansion in the out-of-plane direction.

### Depletion of condensins results in softer chromosomes

We isolated single mitotic chromosomes from WT, control siRNA-treated, and condensin-depleted (SMC2 siRNA) HeLa cells for mechanical experiments. Fig. 3A shows a WT chromosome suspended between two micropipettes. WT chromosomes are homogeneous, and they stretch uniformly under applied force (Fig. 3Ab-c). They display linear, reversible elasticity (Fig. 3Ad), with force applied during extension matching that observed during retraction, for forces of up to 300 pN (linear regime extends to ~300 pN, reversible regime is broader, extending to about three times the doubling force, or to ~800 pN) (Sun et al., 2011).

**Figure 3.**
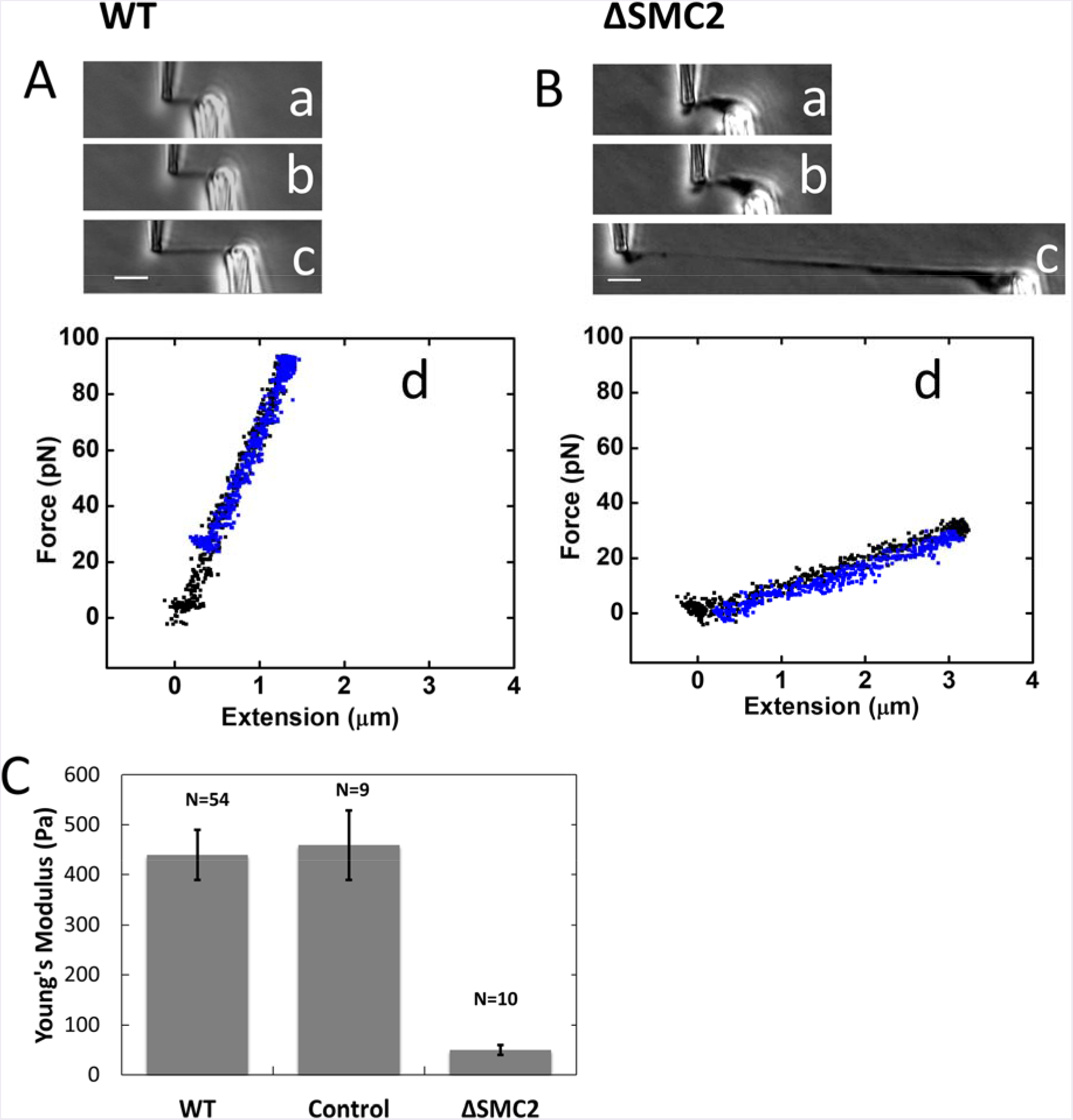
Depletion of condensins results in softening of chromosomes as measured by chromosome mechanics. (A-B) Single chromosomes isolated from untreated (WT) cells (A) and RNAi condensin SMC2-depleted cells (B) were stretched under applied force ((a)-(c)). Initial stretching ((a) and (b)) gave raise to linear reversible force-extension curves (d); extension (black) coincides with subsequent retraction (blue). (c) shows chromosome stretching using greater than 300 pN force, qualitatively showing the uniform stretching of the untreated chromosomes, versus the elastic failure that occurs in the SMC2-depleted case. Bar = 5 μm. (C) Young’s modulus of untreated chromosomes, chromosomes from cells 36 hrs after transfection of non-targeting control siRNA, and chromosomes from cells 36 hrs after transfection of condensin SMC2-targeting siRNA.

By contrast, chromosomes from condensin-depleted (36 hours following SMC2-targeted siRNA treatment) cells showed inhomogeneous compaction, displaying thin and thick regions (Fig. 3Ba). A reversible elastic response of isolated chromosomes was observed over a short range of stretching (Figs. 3Bb, d). However, for applied forces of ~200 pN chromosomes extended in a nonuniform manner, with thinner and thicker regions along their length (Fig. 3Bc). For long extensions, the stretching force generated during retraction was lower than that observed during extension, indicating irreversible elastic response likely isolated with substantial chromosome unfolding (Fig. S3B; note that in addition to irreversibility there was likely some instrumental drift in this long-extension experiment).

From chromosome stretching experiments we obtain an estimate of chromosome elastic modulus (Young’s modulus) (Sun et al., 2011). Young’s modulus measures the force per unit area required to double the length of an elastic object if its initial linear elastic response were extrapolated to that length, and provides a shape-independent quantification of elastic stiffness. We take this measure of elastic stiffness as a quantification of the degree of interconnection of chromatin fibers in the interior of a chromosome (Poirier and Marko, 2002, Sun et al., 2011). Cross-sectional area was obtained optically by measuring the average radius of chromosomes in phase contrast images (Sun et al., 2011). We found the Young’s modulus of WT chromosomes (untreated HeLa cells) to be 440 ± 50 Pa. while the Young’s modulus of condensin-depleted chromosomes (36 hrs after SMC2 siRNA treatment) was found to be 50 ± 10 Pa.

Experiments for chromosomes from cells 36 hours following treatment with non-targeting control siRNA yielded a modulus of 460 ±70 Pa, within our measurement error equal to the result for the WT chromosomes.

### Antibody labeling of condensin shows a discontinuous pattern along mitotic chromosomes

In order to visually examine the condensin distribution on chromosomes, we developed an immunofluorescence staining technique for unfixed, isolated chromosomes. In brief, primary and secondary antibodies were sequentially introduced onto the chromosomes via a third spray pipette with a 5 μm diameter opening. This “microspraying” was carried out for 10 min by applying 10-100 Pa pressure through the spray pipette (Fig. 4A). Primary and secondary antibodies were diluted in 50% phosphate buffered saline (PBS) buffer to increase antibody accessibility (Poirier et al., 2002), and to minimize protein dissociation from chromosomes (lowering salt concentration increases binding affinity of most proteins to DNA by strengthening electrostatic interactions). A washing step between primary and secondary sprays was carried out by moving the isolated chromosome into a different region of the sample cell, far away from the spray location. Following microspraying, flow was stopped for several minutes to allow the small volume of antibody sprayed into solution (~5-20 μl) to diffuse away into the 1.5 ml dish volume. Comparison of phase contrast images before and after spraying indicated that the rod-shaped metaphase chromosome morphology was not disturbed by flow or by binding of antibodies (see, *e.g.*, Fig. 4Ba).

**Figure 4.**
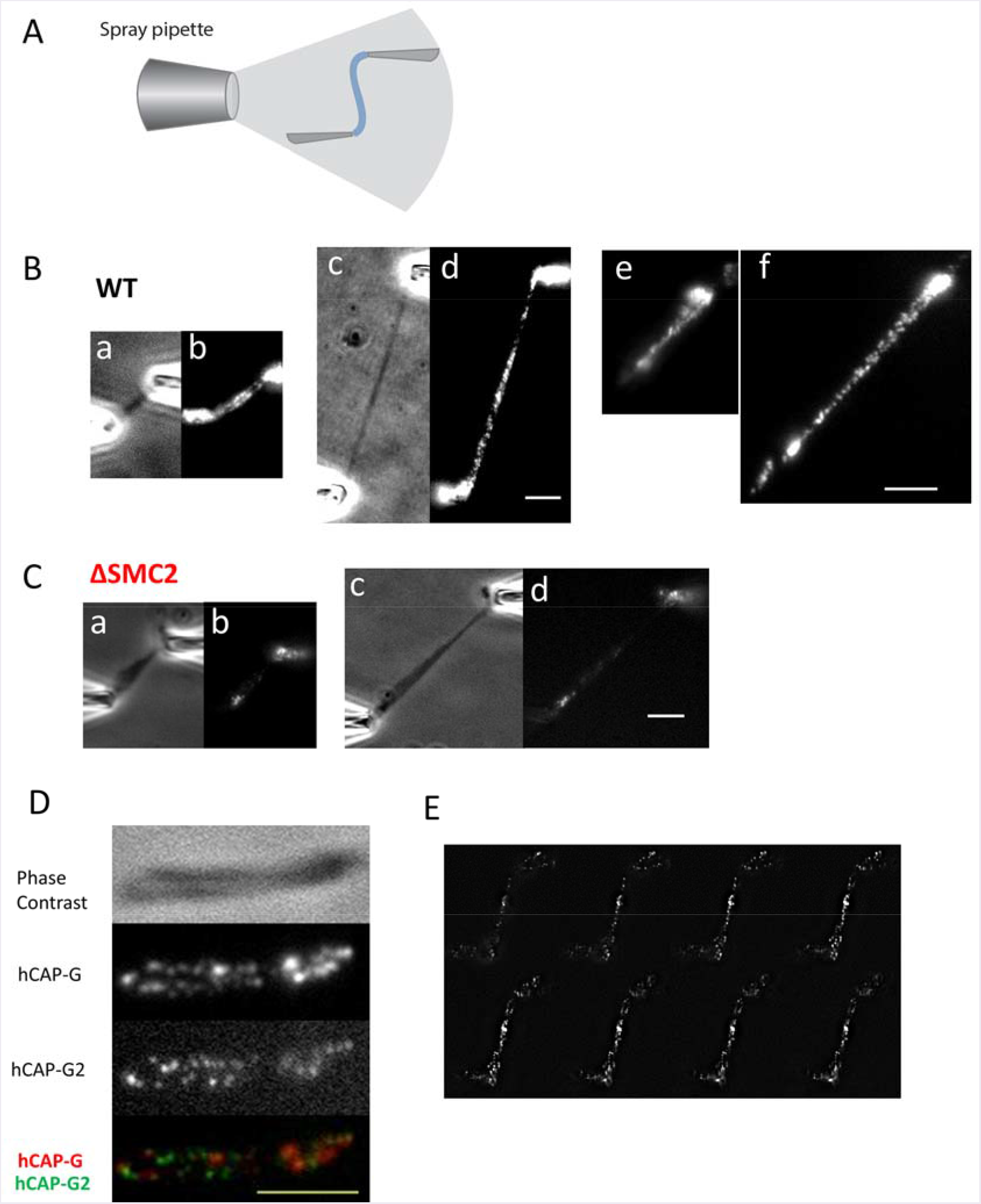
Condensin displays a discontinuous distribution on mitotic chromosomes. (A) Microspray setup for immunofluorescence labeling. Single chromosomes from control (B) and condensin depleted (C) cells, imaged with either phase contrast ((a) and (c)) or stained with anti-SMC2 ((b) and (d)); (c) and (d) show the stretched chromosomes of (a) and (b). Panels (e) and (f) show SMC2 staining results from a separate experiment. (D) A native, 2X stretched chromosome obtained from chromosome isolation by micropipettes, imaged using phase contrast, and anti-hCAP-G and anti-hCAP-G2 immunofluorescence. Merged image is shown in the lowest panel. (E) Representative images of condensin I (anti-hCAP-G) distribution in z-stack from top to bottom, showing a clear dotted pattern. Images were taken 200 nm apart in z axis. Bar = 5μm.

Figs. 4Bb and 4Bd shows the SMC2 staining along an isolated chromosome, indicative of the total condensin distribution, since SMC2 is common to both condensins I and II. Before stretching (Fig. 4Bb), we observed staining along the chromosome, concentrated in the central (centromeric) region, but with staining along the entire length of the chromosome. Although some variation in staining is observed along the chromosome, we observed SMC2 staining along the interior regions of the two adjacent chromatids, similar to the result of prior condensin-antibody-labeling experiments on fixed and spread chromosomes (Ono et al., 2003, Maeshima and Laemmli, 2003, Ono et al., 2004, Lai et al., 2011, Samejima et al., 2012).

Use of moderate tension allows us to stretch out the chromosomes to better examine their structure. Stretching revealed that the SMC2 staining had a discontinuous pattern along the WT chromosome (Fig. 4Bd). A clearly disconnected pattern became clear when chromosomes were stretched to ~ 2 to 3-fold native length, which revealed a clear separation of individual SMC2-containing loci (Fig. 4Bd). Experiments with antibodies diluted in 100% PBS showed less binding overall consistent with the binding-weakening effect of higher salt, but the staining still showed a disconnected SMC2 pattern when the chromosomes were stretched (Fig. S4). Discontinuous patterning was also present when the chromosome was stretched before antibody spraying (Fig. S5A-B). The total fluorescence after extension was observed to be approximately the same as before extension indicating that there was not extensive loss of condensin during the stretching experiments, although we cannot rule out that some condensins may be lost during chromosome extension.

To address the possibility that the discontinuous SMC2 pattern was due to the bidentate nature of full-length antibodies, we carried out the same type of experiment using Fab fragments for both primary (anti-SMC2) and secondary antibody sprays (Fab fragments contain only single antigen binding sites). Following Fab fragment primary and secondary sprays, we again observed a discontinuous pattern of SMC2 along the chromosome (Fig. S5C-D), similar to the result for the full-length antibody experiment.

To examine the possibility that the condensin staining pattern we observed was due to incomplete or selective labeling of a continuous, homogeneous structure, we compared the deformability of the SMC2-antibody-labeled loci with that of the unlabeled regions, as we gradually stretched the chromosomes from their native length to 3-fold their native length. We measured the diameter of each condensin SMC2 cluster, and the distance between neighboring clusters, during stretching (cluster sizes were quantified using line scan analysis along the chromosome, using the full width at half maximum of fluorescence intensity; inter-cluster distances were determined by similar line-scan analysis using the peak-to-peak distance for adjacent clusters). We found that as chromosomes were stretched to three-fold their native lengths, the apparent cluster sizes stayed approximately the same, while the distances between adjacent clusters increased approximately 3-fold (Fig. S6). The cluster sizes we observe are comparable to the expected diffraction-limited size of a point emitter (~250 nm) and therefore we cannot rule out that they are not stretching at all; given their intensities, we are certain that each cluster contains many fluorescently labeled antibodies. These data suggest that the SMC2-rich regions along the chromosome likely were stiffer than the SMC2-poor inter-loci regions, with the SMC2-poor regions generating most of the stretching of the metaphase chromosome. Similar results were obtained from similar experiments using SMC2 antibody diluted in 100% PBS (Fig. S4). This difference in stiffness further suggests that the cluster pattern we observed is not due to selective labeling of a continuous structure.

Fig. 4C shows the SMC2 pattern on a condensin-depleted (SMC2-siRNA-treated) chromosome, using the same antibody staining and imaging procedure as for the WT case. Much less staining was observed, indicating that much less condensin was associated with the chromosome than in the wild-type case. For condensin-depleted chromosomes the SMC2 staining was clustered into a small number of spots unevenly distributed along the chromosome.

### Condensin I and II antibodies label disjoint discontinuous regions

To examine the distribution of condensin I and condensin II staining along WT mitotic chromosomes, we simultaneously labeled condensin I- and condensin II-specific subunits hCAP-G and hCAP-G2 on the same chromosome. Fig. 4D shows the hCAP-G and hCAP-G2 distribution on a stretched chromosome (extended to 1.5 fold of its native length), showing a discontinuous pattern for both hCAP-G and hCAP-G2. A merged image indicates that condensin I and II have distinct patterns along metaphase chromosomes, with condensin I and II spots alternating along the chromosome arms.

To further investigate the condensin distribution, and in particular to examine the possibility that the condensin I-antibody-labeled regions might actually be part of a continuous coil which goes in and out of the focal plane, we carried out Z-stack imaging of a condensin I-labeled chromosome (again moderately stretched to about 1.5 of its native length), taking images every 200 nm (Fig. 4E, Fig. S7). Fig. 4E shows Z-stacked images of the condensin I distribution, demonstrating a clear dotted pattern at any given focal plane. We also found that the two bright condensin I spots in the middle of the chromosome are coincident with the centromere (CREST staining, Supplementary Material Fig. S8). This centromere enrichment was observed for condensin I and condensin II (Fig. S8, 4D-I).

### Depletion of condensin II impacts chromosome mechanics more than depletion of condensin I

To investigate their differential contributions to chromosome mechanics, condensin I or condensin II were depleted in separate experiments, using siRNAs targeting hCAP-G or hCAP-G2. The average degree of protein depletion 60 hours after siRNA transfection was ~ 90 ± 10 %, and 77 ± 14% for hCAP-G and hCAP-G2 respectively, quantified by Western blot for whole-cell extracts (Fig. 5A). Single chromosomes were isolated from condensin I or condensin II-depleted cells 60 hours following siRNA transfection, following the procedure described above.

**Figure 5.**
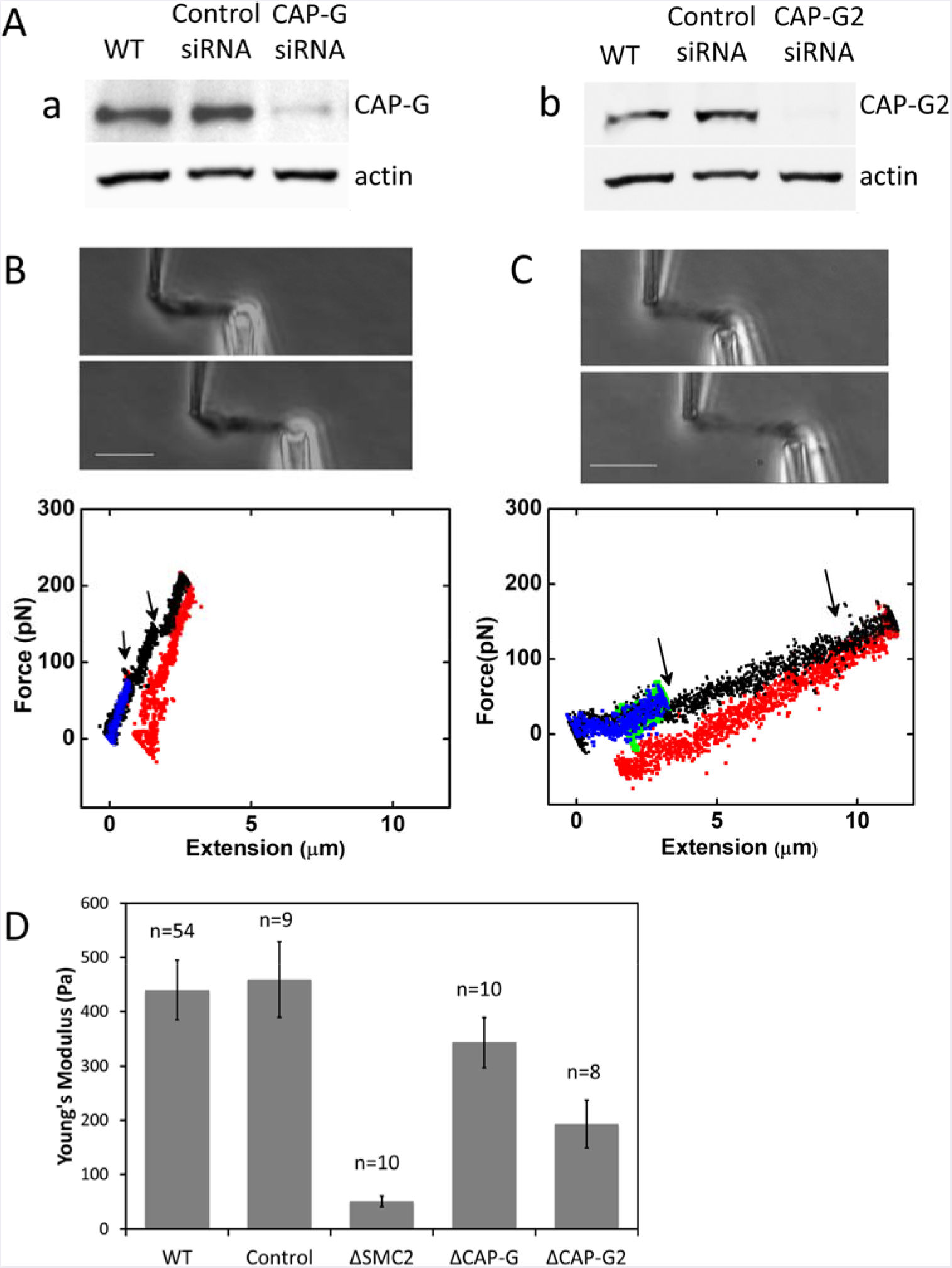
Relative contribution of condensin I and II to chromosome mechanics. (A) RNAi depletion of hCAP-G (a), and hCAP-G2 (b), verified by Western blot. (B-C) Elasticity of condensin I (B) or condensin II depleted chromosomes (C). Top panels: phase contrast images for single chromosome stretching. Bottom panels: Force-extension curves. Short-extension curves (blue and green) are linear and reversible (green return curves are nearly completely obscured by the blue extension curves). Longer extensions (black) show sudden drops in force (arrows), possibly due to opening of locally folded structures. Return curves from long extensions (red) display appreciable irreversibility. (D) Young’s Moduli of chromosomes isolated from condensin I-or condensin II-depleted cells, compared to those of native, control siRNA-treated, and condensin-subunit-depleted chromosomes (SMC2, hCAP-G and hCAP-G2 siRNA-treated).

Knockdown of either condensin I or condensin II had a less severe effect on chromosome mechanics than condensin SMC2 knockdown. The basic morphology of hCAP-G- and hCAP-G2-depleted chromosomes was similar to WT chromosomes (Fig. 5B). The force-extension curves were reversible in the low degree of stretching (<2 fold of native length), but started to show a unique “sawtooth” pattern when they were further extended, suggesting opening up of folded local domains (Fig. 5C: note also the irreversibility of the force response; the slightly negative forces in the figure may be in part due to instrumental drift). Notably, we never observed similar sawtooth opening events for condensin SMC2-depleted chromosomes in this extension range; instead, SMC2-depleted chromosomes simply showed a very soft force response.

The Young’s modulus of hCAP-G-depleted chromosomes is similar to that of WT chromosomes (343 ± 46 Pa, a lower value but within measurement error of the WT result); on the other hand, hCAP-G2-depleted chromosomes are significantly softer than WT chromosomes (193 ± 44 Pa) (Fig. 5D). Since the protein knockdown level is slightly different for condensin I and condensin II, we also linearly corrected the measured Young’s moduli for the partial protein knockdown level measured from Western blot analysis. This gives similar results to those before correction, *i.e.*, a larger effect from condensin II depletion than from condensin I (Fig. S9). Fig. S3 reproduces force-extension curves of Figs. 3 and 5 on the same scales for easy qualitative comparison. We note that while representative, these are force curves from individual experiments, and quantitative comparison of spring constants or moduli requires averaging over multiple experiments.

### Prolonged mitotic arrest by colchicine or nocodazole overcompacts chromosomes, overloading them with condensin and stiffening them

It has been widely recognized that chromosome morphology is altered when cells are stalled in metaphase using mitotic spindle inhibitors such as colchicine and nocodazole. We have observed that metaphase chromosomes from colchicine and nocodazole treated cells are over-compacted compared to native metaphase chromosomes (Fig. 6, A-B), in accord with prior studies (Tomisato Miura, 2012, Foresti et al., 1993, Ostergren, 1944, Rieder and Palazzo, 1992). A reasonable hypothesis is that metaphase chromosomes from WT and metaphase-stalled cells may have appreciable structural differences, and hence differences in chromosome elasticity.

**Figure 6.**
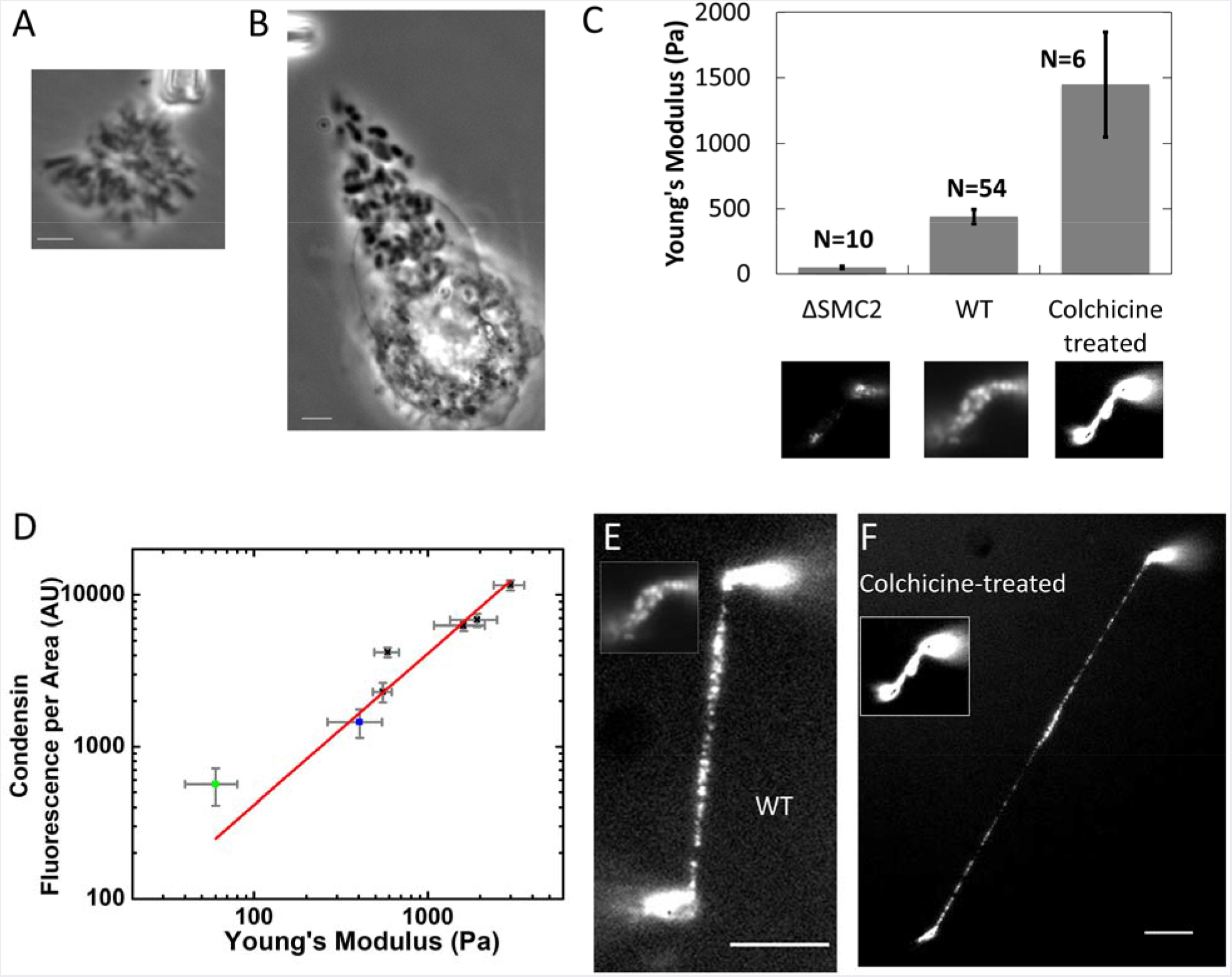
Metaphase-stalled cells have chromosomes which are stiffer and which have more condensin relative to WT. (A-B) phase contrast image of whole genome isolated from untreated (A) and colchicine-treated (B) HeLa cells. (C) Young’s moduli of condensin depleted, native, and colchicine treated chromosomes. Lower panel images show representative examples of condensin immunostaining for each group. (D) Condensin fluorescence per chromosome occupied area versus Young’s modulus for condensin-depleted chromosomes (green squares), native chromosomes (blue squares), and colchicine-treated chromosomes (black squares). (E-F) Side-to-side comparison of native and colchicine-treated chromosome after stretching. Inserts show the chromosomes before stretching. Bar = 5μm. Note that the inset of panels E and F are the same image as the center and lower right insets of panel C, and also that the lower left inset of panel C is the same image shown in Fig. 4Cb.

To investigate the mechanism underlying overly compacted chromosomes in metaphase-blocked cells, we carried out single chromosome micromanipulation and immunofluorescence staining of condensin SMC2 on chromosomes in colchicine-treated cells. The chromosomes from colchicine-treated cells have a much larger Young’s modulus in comparison with those of WT chromosomes (Fig. 6C). We also observed strongly elevated condensin levels in SMC2-antibody labeling experiments on chromosomes isolated from colchicine-blocked cells, relative to the WT case (Fig. 6C, lower panel).

Combining data from colchicine-treated and SMC2 siRNA-treated cells, we plotted condensin fluorescence intensity per unit area on the chromosome as a function of the chromosome Young’s modulus, which showed a linear correlation (Fig. 6D). Condensin-depleted chromosomes from the siRNA experiments have a lower condensin level per unit area, as well as a low elastic modulus (Fig. 6D, green squares), compared with those of native chromosomes (Fig. 6D, blue squares), while chromosomes from colchicine-treated cells at various time points (from 12 hours to 24 hours) have elevated condensin fluorescence per chromosome area, as well as elevated Young’s moduli (Fig. 6D, black squares).

Fig. 6E shows the condensin distribution on native chromosomes versus that of chromosomes from colchicine treated cells on stretched chromosomes. Inserts show the image of the same chromosome before stretching. The condensin distribution on native chromosomes suggests a discontinuous organization even before stretching, and those clusters are more distinct after stretching. For chromosomes from colchicine-treated cells, the condensin distribution appears continuous before stretching, but then displays a similar cluster organization pattern as in the WT case after stretching. We next examined whether the increase in condensin level on chromosomes from colchicine-treated cells is due to an increase in total cellular condensin level. We compared the condensin fluorescence intensity in fixed mitotic cells for untreated and colchicine-treated cells, and found that the overall condensin levels per cell are nearly the same (Fig. S10A). By comparison, the total fluorescence from each chromosome is elevated for chromosomes isolated from colchicine treated cells relative to untreated cells (Fig. S10B). We also examined the condensin I to condensin II ratios on metaphase chromosomes, by simultaneously antibody-labeling hCAP-G and-G2 on colchicine-treated and untreated metaphase chromosomes; the ratio of condensin I to condensin II was not significantly different (Fig. S10C). Finally, in experiments where nocodazole was used to stall the cells in metaphase, overloading of SMC2 onto metaphase chromosomes to similar levels as in in the colchicine experiments was observed (Fig. S11).

## Discussion

### Condensin controls the elastic stiffness of mitotic chromosomes

Our experiments indicate that condensin plays a major role in controlling the elastic stiffness of mitotic chromosomes. WT human chromosomes show robust mechanical elastic response, and can be reversibly extended two-fold by ~300 pN forces comparable to those applied by the mitotic spindle *in vivo* (Nicklas, 1988). The Young modulus of WT chromosomes is ~440 Pa. However, we find that RNAi knockdown of SMC2, common to both condensin I and II, leads to unstable chromosome structure, a strong reduction in chromosome Young’s modulus to ~50 Pa, and irreversible elastic behavior (Figs. 2 and 3). More complete knockdown of SMC2 leads to a complete failure of chromosome compaction and segregation (Fig. 2A). Our results are consistent with earlier qualitative observations of morphological defects (Ono et al., 2003, Hirota et al., 2004) and mechanical instability (Hudson et al., 2003) of condensin-depleted chromosomes.

In addition, we have found that stalling cells in metaphase using spindle-blocking drugs (colchicine, nocodazole) leads to strong (up to roughly threefold) overloading of SMC2, with a concomitant increase in Young’s modulus (again up to approximately threefold). Plotting RNAi and metaphase-stalled data together (Fig. 6D) shows that chromosome stiffness is approximately linearly dependent on SMC2 density on chromosomes as quantified by antibody labeling. We also note that in our experiments we have observed increased amounts of both condensin I and condensin II on chromosomes from metaphase-stalled cells (Fig. S10C), indicating that our metaphase stall was not so long that it led to loss of condensin I as observed in the long-metaphase-stall experiments of (Lai et al., 2011).

### Condensin is organized into discontinuous “centers” along chromosome arms

An advantage of our chromosome manipulation approach is that we are able to carry out quantitative fluorescence microscopy on individual unfixed, unspread, mitotic chromosomes, aligned precisely in the focal plane of a high-numerical-aperture microscope objective. Because of their robust elasticity we can slightly stretch out single metaphase chromosomes, boosting our ability to observe small-scale structural details. Using primary and secondary antibodies directly sprayed onto chromosomes, we have observed discontinuous distributions of SMC2, hCAP-G, and hCAP- G2 along stretched chromosome arms (Fig. 4B,D,E). However, for unstretched chromosomes, the condensin-rich spots come close to one another, appearing to be nearly continuously distributed along the chromatids as seen in prior studies (Ono et al., 2004, Ono et al., 2003). Notably, some prior studies have concluded that the condensin distribution along chromatids is disconnected (Samejima et al., 2012).

The discontinuous labeling distribution suggests that condensin in metaphase chromosomes is organized into localized “condensin centers”. Stretching chromosomes causes little or no distortion of the condensin centers, while generating clear separation of those centers: most of the elastic deformation of chromosomes is due to extension of inter-condensin-center chromatin (Fig. 4). This suggests that the elastic modulus of the highly labeled condensin centers is much larger than that of the inter-condensin-cluster chromatin, and that the condensin-antibody-stained puncta and the unstained regions have different structures. Our results are consistent with a very recent super-resolution imaging study (Walther et al., 2018) where condensins are observed to be separated from one another but with modulation of their density on longer length scales, consistent with the structure seen in our experiment where we have ~250 nm diffraction-limited wide-field imaging resolution.

The interpretation that condensin centers are connected by chromatin with a low concentration of condensin is also consistent with observations that intermittent cutting of DNA alone leads to rapid loss of chromosome elasticity followed by complete chromosome dissolution, essentially converting the WT “elastic solid” chromosome to a “liquid droplet” of chromatin fragments (Poirier and Marko, 2002, Sun et al., 2011). If the condensin centers formed a solid “scaffold” structure inside the metaphase chromosome, one would expect the surrounding chromatin to be cut away, leaving the scaffold behind. The direct observation of condensin centers strongly suggests that the interior of the metaphase chromosome should be considered to be a “gel” or “network” of chromatin, with a structure stabilized by isolated “crosslinking” elements.

The approximate proportionality of chromosome Young’s modulus to the amount of condensin present on the chromosome inferred from SMC2 antibody staining (Fig. 6D) provides further evidence for this “network” model with condensin acting as a crosslinking element, since the elastic modulus of polymer networks is in general proportional to the density of crosslinkers (De Gennes, 1979). A recent study using chromosome conformation capture of mitotic chromosomes was found to be consistent with a model of consecutive chromatin loops stabilized by chromosome organizing protein clusters (Naumova et al., 2013); our results suggest that those chromatin loops are organized into distinct clusters.

It is possible that our antibody-visualization approach underestimates the amount of condensin on metaphase chromosomes, either because antibodies fail to have access to all their targets in the native chromosome, or perhaps because the antibodies do not bind sufficiently tightly to be stable on the unfixed chromosomes in the extracellular medium. However, even if this is the case, the difference in elasticity of the condensin centers versus the adjacent chromatin indicates that condensin is inhomogeneously distributed along metaphase chromosomes. Studies of cells carrying bright, functional, gfp-fusion condensing subunits could strengthen the conclusion that condensin is organized discontinuously along metaphase chromosomes.

### Differential contribution of condensin I and II to mitotic chromosome structure and stiffness

Condensin I and II have been proposed to perform non-overlapping functions in mitotic chromosome organization (Ono et al., 2003, Hirano, 2012). Consistent with this we have observed distinct and alternating condensin I and condensin II centers along chromosome arms, the alternating pattern of I and II in agreement with prior studies (Ono et al., 2004, Ono et al., 2003). Condensin I- and II-rich centers (as visualized by antibodies to hCAP-G and -G2) are distinct, with roughly 20 condensin I and 20 condensin II centers along the ~150 Mb chromosomes we have studied (Fig. 4). We estimate that there is one condensin center for every ~5 Mb of DNA, and also that each condensin center involves many (roughly 50) condensin complexes, since current estimates are that there is approximately one condensin per 90 kb of DNA in metazoan chromosomes (Samejima et al., 2012). The apparent clustering of condensins in metaphase chromosomes may be indicative of cooperative interactions between individual condensin complexes. Even without any cooperative interaction between SMCs, “loop-extrusion” dynamics could drive condensin accumulation at the bases of chromatin loop domains while compacting chromosomes (Alipour and Marko, 2012, Goloborodko et al., 2016b, Goloborodko et al., 2016a). Single-molecule experiments have indeed observed DNA translocation (Terakawa et al., 2017) and loop-extrusion (Ganji et al., 2018) dynamics for yeast condensin, supporting this model. A possible molecular mechanism underlying loop extrusion by condensin which is consistent with the observations of (Ganji et al., 2018) has been presented in (Lawrimore et al., 2017).

We find that the centromere has a SMC2-rich center associated with it, in accord with recent observations in yeast (Stephens et al., 2013). We further find the centromeric condensin center to be enriched in condensin I and condensin II, rather than only condensin II as observed by (Ono et al., 2004). The reason for this difference is unclear, but it may reflect the different conditions for the antibody binding, *i.e.*, the native solvated chromosomes of this study *vs.* fixed chromosome spreads used in other studies.

We have found that the stiffness of the metaphase chromosome is more dependent on condensin II than on condensin I. Condensin I-depleted chromosomes have an average stiffness only slightly less than that of WT chromosomes, while condensin II-depleted cells have significantly softer chromosomes than WT. However, neither condensin I nor condensin II depletion alone has as severe a softening effect as does depletion of both condensin complexes at once (via SMC2 knockdown).

Our results support the hypothesis that condensin I and condensin II are involved in different levels of chromosome compaction. This is consistent with studies showing that condensin II associates with chromosomes in the nucleus to generate thin prophase chromosomes, while condensin I binds chromosomes only after nuclear envelope breakdown, driving further chromosome compaction (Ono et al., 2003, Hirota et al., 2004). Our results suggest that condensin II serves as a major chromatin crosslinking element which determines the stiffness of mitotic chromosomes, while condensin I drives longitudinal compaction, compacting chromosomes without significantly increasing the crosslinking density (and therefore the elastic modulus). The morphology of condensin I or II-depleted chromosomes remains the basic metaphase rod shape. The difference between the effects of condensin I and II knockdowns is also consistent with the dynamics of the two complexes during metaphase (when our isolations are done), namely that condensin II is found to be essentially immobile while condensin I is highly dynamic (Hirota et al., 2004).

Interestingly, the stretching curves of isolated chromosomes from condensin I or condensin II depleted cells displayed a “sawtooth” pattern, with sudden drops in applied force, indicating local domain opening events (Fig. 5). We did not observe this type of stretching pattern with either native chromosomes or condensin/SMC2-depleted chromosomes. The condensin/SMC2-depleted chromosomes may be simply too soft to detect such “jumps”.

### Condensin drives removal of inter-chromosome entanglements

We have observed severe defects in the individualization of chromosomes in condensin-depleted cells. This becomes most dramatic in later stages of condensin depletion, where it appears that all the chromosomes in the cell become one massive entangled globule of chromatin, with no sign of chromosome individualization or sister chromatid resolution (Fig 2A). This is likely related to the appearance of anaphase chromatin bridges in condensin defective cells (Saka et al., 1994, Sutani et al., 1999) which reflects a failure to resolve entanglements between sister chromatids. These results suggest that absence of condensin leads to a general failure of removal of both intra- and inter-chromosome entanglements.

Our experiments with cells stalled in metaphase show the opposite effect: we observe highly compacted, condensin-overloaded, mechanically stiff, and well individualized chromosomes. For many years it has been observed that chromosomes in metaphase-stalled cells are more scattered over the cell and are hypercompacted (Ostergren, 1944). Our results indicate that this behavior is correlated with condensin over-loading; removal of interchromosome entanglements, like Young’s modulus, is essentially dependent on condensin level on chromosomes: the more condensin on a chromosome, the more complete is the entanglement removal process.

We have observed that interchromosome linkers, which are present at WT metaphase (Maniotis et al., 1997, Marko, 2008), are nearly absent in metaphase-stalled cells, possibly indicative of a more complete chromosome individualization process. The interchromosome linkers may be chromatin regions which are particularly resistant to entanglement removal, but which are eventually disentangled given a large enough amount of chromosomal condensin. Alternately, the linkers may be cut in metaphase-stalled cells by DNA-cleaving mechanisms known to be active in cells held in metaphase by microtubule-blocking drugs (Ganem and Pellman, 2012, Orth et al., 2012).

### Condensin-depleted chromosomes can fold, but are unstable

Intriguingly, there is a dramatic difference in stability of chromosomes from condensin-depleted cells compared with those of WT cells. WT chromosomes are very stable, while condensin-depleted chromosomes fall apart after extraction (Fig. 2B). There is still residual condensin in condensin-depleted cells (Fig. 4C), since RNAi leads to progressive dilution of the amount of condensin after each cell cycle. Thus, the change in chromosome stability may indicate that formation of stable condensin clusters in native chromosomes is cooperative; at low condensin concentration, the isolated condensin complexes may simply fall apart after chromosome extraction into physiological buffer. This could be tested using gfp-labeled condensins, via direct observation of the amount of labeling with time following chromosome extraction.

Alternatively, it may be the case that other proteins that play a major role in metaphase chromosome folding require the presence of condensins for their stable binding and therefore for stability of metaphase chromosomes. As an example, it has been suggested that in addition to condensins, there are “regulators of chromosome architecture”, additional factors which act to stabilize metaphase chromosome folding (Vagnarelli et al., 2006, Samejima et al., 2012). We note that conventional chromosome isolation/preparation protocols which include hypotonic buffers and washing steps, when applied to chromosomes from SMC2-siRNA-treated cells, would likely lead to appreciable chromosome unfolding of the sort seen in Fig 2Bd.

### Condensin loading and chromosome compaction are kinetically controlled

Our experiments with cells stalled in metaphase indicate that given enough time spent in metaphase, condensin will “overload” onto metaphase chromosomes, with additional condensin from the cellular pool apparently binding and driving stronger compaction, as more time is spent in metaphase. Interestingly, we did not detect a significant change in condensin I to condensin II ratios in metaphase-stalled cells, indicating that both condensin I and condensin II are overloaded in an extended metaphase. This suggests chromosome compaction is a dynamic process, with normal cells terminating chromosome compaction well before the point of maximal condensin loading, compaction, and chromosome/chromatid separation. The regulatory mechanism for condensin loading is not understood, but a simple possibility consistent with our results is that condensin is simply progressively loaded throughout mitosis, with overloading occurring when metaphase is artificially extended. Alternately condensin may be overloaded in response to chromosomal damage that occurs in a prolonged metaphase, *e.g.*, dsDNA breaks (Orth et al., 2012), since condensin has been implicated in prevention of accumulation of DNA damage (Sakamoto et al., 2011, Schuster et al., 2013). We do note that it is possible that there is some condensin I loss during isolation of metaphase chromosomes, given its dynamic binding during mitosis (Gerlich et al., 2006).

### Condensin function: chromosome folding and chromatid disentanglement

Our experiments are consistent with the notion that metaphase chromosomes can be considered as “chromatin gels” Poirier and Marko, 2002), in which condensin complexes (or clusters of them) play the role of isolated chromatin-cross-linking elements. However, if condensin were to crosslink chromatin indiscriminately, the result would be generation of entanglements by topo II rather than removal of them. (Marko and Siggia, 1997, Marko, 2011). In order to avoid this, condensin may act as a mediator of *lengthwise compaction*, acting *in cis* along chromatin to fold up chromosomes without linking separate DNA molecules together.

A conceivable mechanism for lengthwise compaction is progressive “extrusion” of chromatin loops by condensins (Marko, 2011, Alipour and Marko, 2012, Marko, 2009, Nasmyth, 2001, Goloborodko et al., 2016a, Goloborodko et al., 2016b, Gibcus et al., 2018, Ganji et al., 2018). In such a model, condensin II would act to push chromatin to the exterior of the chromosome, perhaps in concert with condensin-associated factors such as the chromokinesin KIF4, (Samejima et al., 2012), with crowding of chromatin loops providing a thermodynamic drive for topo II to remove interlocks between different chromatids and chromosome. This cooperation between condensin and topo II would lead to separated chromatids and chromosomes (Bhat et al., 1996) with condensin clusters at the loop bases in the interior of chromatids, consistent with the pattern of isolated condensin clusters that we have observed. In the absence of condensin, topo II can be expected to make undirected strand passages, and to globally entangle chromosomes, as we have observed (Fig. 2D).

Despite direct observation of loop-extrusion dynamics by yeast cohesin (Ganji et al., 2018), it remains conceivable that simpler chromatin-looping mechanisms may dominate metaphase chromosome self-organization (Cheng et al., 2015, Marko and Siggia, 1997). However, whatever the detailed molecular mechanisms, our observations are in excellent accord with the general model that condensin and topoisomerase II work together to segregate chromosomes, with condensin compacting chromatin and topo II changing chromatin topology (Hirano, 1995). In this model, condensin drives lengthwise compaction and chromosome stiffening, converting chromosome entanglement into osmotic and mechanical stresses; release of those stresses by topoisomerase II then individualizes chromosomes and resolves sister chromatids (Fig. 2B) (Marko, 2008, Marko, 2009, Marko, 2011, Marko and Siggia, 1997, Hirano, 1995).

**List of Abbreviations**
(h)CAP: (human) Chromosome-Associated Protein
SMC: Structural Maintenance of Chromosomes (complex)
siRNA: small interfering RNA
WT: Wild-Type (untreated)
NEB: Nuclear Envelope Breakdown
CREST: Calcinosis, Raynaud’s phenomenon, Esophageal dysmotility, Sclerodactyly, and Telangiectasia (syndrome)
CCD: Charge-Coupled Device
PBS: Phosphate-Buffered Saline
DMEM: Delbecco Modified Eagle Medium
FBS: Fetal Bovine Serum
Pa: Pascals (SI unit of pressure)
N: Newton (SI unit of force)

## Acknowledgements

This work was supported by the NIH through grants R01-GM105847, U54-CA193419 (CR-PS-OC) and a subcontract to grant U54-DK107980, and by the NSF through grants MCB-1022117, DMR-1611076 and DMR-1206868.

## Author Contribution Statement

MS, RB, JH and JFM conceived and designed the research. MS, RB and JH conducted experiments. MS, RB, JH and JFM analyzed data. MS, RB, JH and JFM wrote, read and approved the manuscript.

## References

Adolphs KW, Cheng SM, Paulson JR, Laemmli UK (1977) Isolation of a protein scaffold from mitotic HeLa cell chromosomes. Proc Natl Acad Sci U S A, 74:4937-4941.

Alipour E, Marko JF (2012) Self-organization of domain structures by DNA-loop-extruding enzymes. Nucleic Acids Res, 40:11202-11212.

Bak AL, Zeuthen J, Crick FH (1977) Higher-order structure of human mitotic chromosomes. Proc Natl Acad Sci U S A, 74:1595-1599.

Bhat MA, Philp AV, Glover DM, Bellen HJ (1996) Chromatid segregation at anaphase requires the barren product, a novel chromosome-associated protein that interacts with Topoisomerase II. Cell, 87:1103-1114.

Cheng TM, Heeger S, Chaleil RA et al. (2015) A simple biophysical model emulates budding yeast chromosome condensation. Elife, 4:e05565.

De Gennes PG (1979) Scaling concepts in polymer physics. Cornell university press, Section V.

Earnshaw WC, Laemmli UK (1983) Architecture of metaphase chromosomes and chromosome scaffolds. J Cell Biol, 96:84-93.

Foresti F, Oliveira C, Dealmeidatoledo LF (1993) A Method for Chromosome Preparations from Large Fish Specimens Using in-Vitro Short-Term Treatment with Colchicine. Experientia, 49:810-813.

Ganem NJ, Pellman D (2012) Linking abnormal mitosis to the acquisition of DNA damage. J Cell Biol, 199:871-881.

Ganji M, Shaltiel IA, Bisht S et al. (2018) Real-time imaging of DNA loop extrusion by condensin. Science, 360:102-105.

Gerlich D, Hirota T, Koch B, Peters JM, Ellenberg J (2006) Condensin I stabilizes chromosomes mechanically through a dynamic interaction in live cells. Curr Biol, 16:333-344.

Gibcus JH, Samejima K, Goloborodko A et al. (2018) A pathway for mitotic chromosome formation. Science, 359.

Goloborodko A, Imakaev MV, Marko JF, Mirny L (2016a) Compaction and segregation of sister chromatids via active loop extrusion. Elife, 5.

Goloborodko A, Marko JF, Mirny LA (2016b) Chromosome Compaction by Active Loop Extrusion. Biophys J, 110:2162-2168.

Green LC, Kalitsis P, Chang TM et al. (2012) Contrasting roles of condensin I and condensin II in mitotic chromosome formation. J Cell Sci, 125:1591-1604.

Hirano T (1995) Biochemical and genetic dissection of mitotic chromosome condensation. Trends Biochem Sci, 20:357-361.

Hirano T (2006) At the heart of the chromosome: SMC proteins in action. Nat Rev Mol Cell Biol, 7:311-322.

Hirano T (2012) Condensins: universal organizers of chromosomes with diverse functions. Genes Dev, 26:1659-1678.

Hirano T, Mitchison TJ (1994) A heterodimeric coiled-coil protein required for mitotic chromosome condensation in vitro. Cell, 79:449-458.

Hirota T, Gerlich D, Koch B, Ellenberg J, Peters JM (2004) Distinct functions of condensin I and II in mitotic chromosome assembly. J Cell Sci, 117:6435-6445.

Hoskins GC (1968) Sensitivity of micrurgically removed chromosomal spindle fibres to enzyme disruption. Nature, 217:748-750.

Hudson DF, Vagnarelli P, Gassmann R, Earnshaw WC (2003) Condensin is required for nonhistone protein assembly and structural integrity of vertebrate mitotic chromosomes. Dev Cell, 5:323-336.

Lai SK, Wong CH, Lee YP, Li HY (2011) Caspase-3-mediated degradation of condensin Cap-H regulates mitotic cell death. Cell Death Differ, 18:996-1004.

Lawrimore J, Friedman B, Doshi A, Bloom K (2017) RotoStep: A Chromosome Dynamics Simulator Reveals Mechanisms of Loop Extrusion. Cold Spring Harb Symp Quant Biol, 82:101-109.

Maeshima K, Laemmli UK (2003) A two-step scaffolding model for mitotic chromosome assembly. Dev Cell, 4:467-480.

Maniotis AJ, Bojanowski K, Ingber DE (1997) Mechanical continuity and reversible chromosome disassembly within intact genomes removed from living cells. J Cell Biochem, 65:114-130.

Marko JF (2008) Micromechanical studies of mitotic chromosomes. Chromosome Res, 16:469-497.

Marko JF (2009) Linking topology of tethered polymer rings with applications to chromosome segregation and estimation of the knotting length. Phys Rev E Stat Nonlin Soft Matter Phys, 79:051905.

Marko JF (2011) Scaling of Linking and Writhing Numbers for Spherically Confined and Topologically Equilibrated Flexible Polymers. J Stat Phys, 142:1353-1370.

Marko JF, Siggia ED (1997) Polymer models of meiotic and mitotic chromosomes. Mol Biol Cell, 8:2217-2231.

Nasmyth K (2001) Disseminating the genome: joining, resolving, and separating sister chromatids during mitosis and meiosis. Annu Rev Genet, 35:673-745.

Naumova N, Imakaev M, Fudenberg G et al. (2013) Organization of the mitotic chromosome. Science, 342:948-953.

Nicklas RB (1988) The forces that move chromosomes in mitosis. Annu Rev Biophys Biophys Chem, 17:431-449.

Ono T, Fang Y, Spector DL, Hirano T (2004) Spatial and temporal regulation of condensins I and II in mitotic chromosome assembly in human cells. Mol Biol Cell, 15:3296-3308.

Ono T, Losada A, Hirano M, Myers MP, Neuwald AF, Hirano T (2003) Differential contributions of condensin I and condensin II to mitotic chromosome architecture in vertebrate cells. Cell, 115:109-121.

Ono T, Sakamoto C, Nakao M, Saitoh N, Hirano T (2017) Condensin II plays an essential role in reversible assembly of mitotic chromosomes in situ. Mol Biol Cell, 28:2875–2886.

Orth JD, Loewer A, Lahav G, Mitchison TJ (2012) Prolonged mitotic arrest triggers partial activation of apoptosis, resulting in DNA damage and p53 induction. Mol Biol Cell, 23:567-576.

Ostergren G (1944) Colchicine mitosis, chromosome contraction, narcosis and protein chain folding. Hereditas, 30:429-467.

Poirier M, Eroglu S, Chatenay D, Marko JF (2000) Reversible and irreversible unfolding of mitotic newt chromosomes by applied force. Mol Biol Cell, 11:269-276.

Poirier MG, Marko JF (2002) Mitotic chromosomes are chromatin networks without a mechanically contiguous protein scaffold. Proc Natl Acad Sci U S A, 99:15393-15397.

Poirier MG, Monhait T, Marko JF (2002) Reversible hypercondensation and decondensation of mitotic chromosomes studied using combined chemical-micromechanical techniques. J Cell Biochem, 85:422-434.

Pope LH, Xiong C, Marko JF (2006) Proteolysis of mitotic chromosomes induces gradual and anisotropic decondensation correlated with a reduction of elastic modulus and structural sensitivity to rarely cutting restriction enzymes. Mol Biol Cell, 17:104-113.

Rieder CL, Palazzo RE (1992) Colcemid and the mitotic cycle. J Cell Sci, 102(Pt 3):387-392.

Saka Y, Sutani T, Yamashita Y et al. (1994) Fission yeast cut3 and cut14, members of a ubiquitous protein family, are required for chromosome condensation and segregation in mitosis. EMBO J, 13:4938-4952.

Sakamoto T, Inui YT, Uraguchi S et al. (2011) Condensin II alleviates DNA damage and is essential for tolerance of boron overload stress in Arabidopsis. Plant Cell, 23:3533-3546.

Samejima K, Samejima I, Vagnarelli P et al. (2012) Mitotic chromosomes are compacted laterally by KIF4 and condensin and axially by topoisomerase IIalpha. J Cell Biol, 199:755-770.

Schindelin J, Arganda-Carreras I, Frise E et al. (2012) Fiji: an open-source platform for biological-image analysis. Nat Methods, 9:676-682.

Schuster AT, Sarvepalli K, Murphy EA, Longworth MS (2013) Condensin II subunit dCAP-D3 restricts retrotransposon mobilization in Drosophila somatic cells. PLoS Genet, 9:e1003879.

Somma MP, Fasulo B, Siriaco G, Cenci G (2003) Chromosome condensation defects in barren RNA-interfered Drosophila cells. Genetics, 165:1607-1611.

Steffensen S, Coelho PA, Cobbe N et al. (2001) A role for Drosophila SMC4 in the resolution of sister chromatids in mitosis. Curr Biol, 11:295-307.

Stephens AD, Quammen CW, Chang B, Haase J, Taylor RM, 2nd, Bloom K (2013) The spatial segregation of pericentric cohesin and condensin in the mitotic spindle. Mol Biol Cell.

Strunnikov AV, Hogan E, Koshland D (1995) SMC2, a Saccharomyces cerevisiae gene essential for chromosome segregation and condensation, defines a subgroup within the SMC family. Genes Dev, 9:587-599.

Sun M, Kawamura R, Marko JF (2011) Micromechanics of human mitotic chromosomes. Phys Biol, 8:015003.

Sutani T, Yuasa T, Tomonaga T, Dohmae N, Takio K, Yanagida M (1999) Fission yeast condensin complex: essential roles of non-SMC subunits for condensation and Cdc2 phosphorylation of Cut3/SMC4. Genes Dev, 13:2271-2283.

Terakawa T, Bisht S, Eeftens JM, Dekker C, Haering CH, Greene EC (2017) The condensin complex is a mechanochemical motor that translocates along DNA. Science, 358:672-676.

Tomisato Miura AN, Kosuke Kasai, Mitsuaki Yoshida (2012) Effects of Colcemid-Block on Chromosome Condensation in Metaphase Analysis and Premature Chromosome Condensation Assays. Radiation Emergency Medicine, 1:5.

Vagnarelli P, Hudson DF, Ribeiro SA et al. (2006) Condensin and Repo-Man-PP1 co-operate in the regulation of chromosome architecture during mitosis. Nat Cell Biol, 8:1133-1142.

Walther N, Hossain MJ, Politi AZ et al. (2018) A quantitative map of human Condensins provides new insights into mitotic chromosome architecture. J Cell Biol, 217:2309-2328.

Yeong FM, Hombauer H, Wendt KS et al. (2003) Identification of a subunit of a novel Kleisin-beta/SMC complex as a potential substrate of protein phosphatase 2A. Curr Biol, 13:2058-2064.

